# A unified framework for perceived magnitude and dicriminability of sensory stimuli

**DOI:** 10.1101/2022.04.30.490146

**Authors:** Jingyang Zhou, Lyndon R. Duong, Eero P. Simoncelli

## Abstract

The perception of sensory attributes is often quantified through measurements of sensitivity (the ability to detect small stimulus changes), as well as through direct judgements of appearance or intensity. Despite their ubiquity, the relationship between these two measurements remains controversial and unresolved. Here, we propose a framework in which they arise from different aspects of a common representation. Specifically, we assume that judgements of stimulus intensity (e.g., as measured through rating scales) reflect the mean value of an internal representation, and sensitivity reflects a combination of mean value and noise properties, as quantified by the statistical measure of Fisher Information. Unique identification of these internal representation properties can be achieved by combining measurements of sensitivity and judgments of intensity. As a central example, we show that Weber’s law of perceptual sensitivity can co-exist with Stevens’ power-law scaling of intensity ratings (for all exponents), when the noise amplitude increases in proportion to the representational mean. We then extend this result beyond the Weber’s law range by incorporating a more general and physiology-inspired form of noise, and show that the combination of noise properties and sensitivity measurements accurately predicts intensity ratings across a variety of sensory modalities and attributes. Our framework unifies two primary perceptual measurements – thresholds for sensitivity and rating scales for intensity – and provides a neural interpretation for the underlying representation.

**Significance Statement:** Perceptual measurements of sensitivity to stimulus changes and stimulus appearance (intensity) are ubiquitous in the study of perception. However, the relationship between these two seemingly disparate measurements remains unclear. Proposals for unification have been made for over 60 years, but they generally lack support from perceptual or physiological measurements. Here, we provide a framework that offers a unified interpretation of perceptual sensitivity and intensity measurements, and we demonstrate its consistency with experimental measurements across multiple perceptual domains.

## Introduction

On a blistering summer’s day, we sense the heat. And just as readily, we sense the cooling relief from the onset of a soft breeze. Our ability to gauge the absolute strength of sensations, as well as our sensitivity to changes in their strength, are ubiquitous and automatic. These two judgements have also shaped the foundations of our knowledge of sensory perception.

Perceptual capabilities arise from our internal representations of sensory inputs. Measurements of sensitivity to changes in these inputs have sculpted our understanding of sensory representations across different domains. For example, in the late 1800’s, Fechner proposed that sensitivity to a small change in a stimulus is proportional to the resulting change in the internal representation of that stimulus [1]. By the 1950s, Signal Detection theory was formulated to describe this in terms of stochastic internal representations (e.g. [2, 3]), generalizing beyond Fechner’s implicit assumption that stimuli are represented deterministically. In addition to sensitivity to stimulus changes, humans and animals can also make absolute judgements of stimulus intensities [4–8]. But the experimental methods by which this can be quantified are more controversial [9, 10], and the measurements have proven difficult to relate to sensitivity measurements [11–14].

Consider the well-known example of Weber’s law, which states that perceptual thresholds for reliable stimulus discrimination scale proportionally with stimulus intensity (equivalently, sensitivity scales inversely with intensity). Weber’s law holds for an impressive variety of stimulus attributes. Fechner’s broadly accepted explanation is that sensitivity reflects the change in an internal representation that arises from a small change in the stimulus (specifically, it reflects the derivative of the function that maps stimulus intensity to representation). For Weber’s law, this implies a logarithmic internal representation. The search for physiological evidence supporting Fechner’s proposal has been ongoing for more than a century, but remains inconclusive (e.g. [4, 15]). In the 1950s, Stevens and others found that human ratings of perceived intensity of a variety of sensory attributes (proposed as an alternative measure of internal representation) follows a power law, with exponents ranging from strongly compressive to strongly expansive [16, 17]. Stevens presented this as a direct refutation of Fechner’s logarithmic hypothesis [11], but offered no means of reconciling the two. Subsequent explanations have generally proposed either that intensity and sensitivity judgements arise from different perceptual representations [18–21], or that the two perceptual tasks involve different nonlinear cognitive transformations [22, 23].

Here, we generalize Fechner’s solution, developing a framework to interpret and unify perceptual sensitivity and intensity judgements of continuous sensory attributes. Specifically, we use a simplified form of Fisher Information to generalize classical Signal Detection theory, and use this to quantify the relationship between perceptual sensitivity and the noisy internal representation. We show that a family of internal representations with markedly different noise properties are all consistent with Weber’s law, but only one form is also consistent with power law intensity percepts. Finally, by incorporating a noise model that is compatible with physiology, we demonstrate that the framework can unify sensitivity and intensity measurements beyond the regime over which Weber’s law and Stevens’ power law hold, and for a diverse set of sensory attributes.

## Results

What is the relationship between perceptual sensitivity, and the internal representations from which it arises? Intuitively, a change in stimulus value (e.g. contrast of an image) leads to a change in internal response. When the change in internal response is larger than the noise variability in that response, we are able to detect the stimulus change. This conceptualization, based on Fechner’s original proposals [1] and formalized in the development of Signal Detection theory in the middle of the 20th century, has provided a successful quantitative framework to analyze and interpret perceptual data [2, 3, 24]. Despite this success, Signal Detection theory formulations are usually not explicit about the transformation of stimuli to internal representations, and most examples in the literature assume that internal responses are corrupted by noise that is additive, independent and Gaussian.

A more explicit relationship between sensitivity and internal representation may be expressed using a statistical tool known as Fisher Information (FI). Specifically, the noisy internal responses (*r*) to a stimulus (*s*) are described by a conditional probability *p*(*r*|*s*), and Fisher Information is defined in terms of a second-order expansion of this probability: *F* (*s*) = 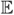[(∂ log *p*(*r*|*s*)*/*∂*s*)^2^]. This quantity specifies the precision with which the stimulus can be recovered from the noisy responses, and 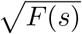 provides a measure of sensitivity to stimulus changes (see Methods). Fisher Information is quite general: it can be used with any continuous stimulus attribute, and any type of response distribution (including multi-modal, discrete, and multi-dimensional responses), although only a subset of cases yield an analytic closed-form expression. In engineering, it is used to compute the minimum achievable error in recovering signals from noisy measurements (known as the “Cramér-Rao bound”). In perceptual neuroscience, it has been used to describe the precision of sensory attributes represented by noisy neural responses [25–28], as a bound on discrimination thresholds [29–31], and to synthesize optimally discriminable stimuli [32].

### Interpreting Weber’s law using Fisher Information

Typically, Fisher Information is used to characterize decoding errors based on specification of an encoder. Here, we are interested in the reverse: we want to constrain properties of an internal representation (an encoder) based on external measurements of perceptual sensitivity (decoder errors). Consider Weber’s law, in which perceptual sensitivity of a stimulus attribute is inversely proportional to the value of the attribute. If we assume obervers achieve the bound expressed by the Fisher Information, this implies that 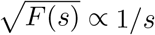. What internal representation, *p*(*r*|*s*), underlies this observation? The answer is not unique. Although the complete family of solutions is not readily expressed, we can deduce and verify a set of three illustrative examples (Fig. 1).

**Figure 1.**
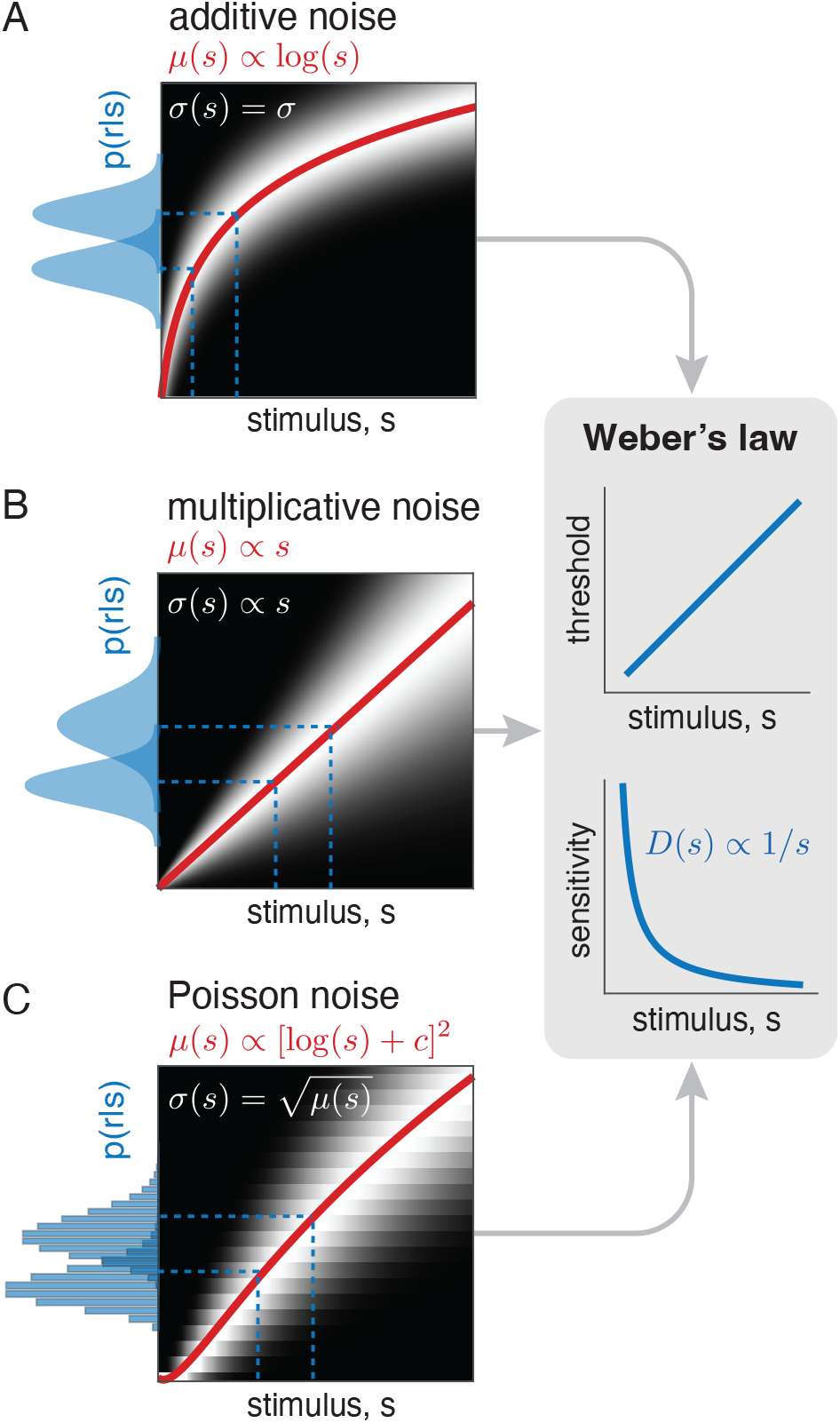
Three different internal representations, each consistent with Weber’s law. Each panel on the left shows a stimulus-conditional response distribution, *p*(*r*|*s*) (grayscale image, brightness proportional to conditional probability), the mean response *µ*(*s*) (red line), and response distributions for two example stimuli (blue, plotted vertically). **A**. Mean response proportional to log(*s*), contaminated with additive Gaussian noise, with constant standard deviation, *σ*(*s*) = *σ*. **B**. Mean response proportional to *s*, with “multiplicative” Gaussian noise (standard deviation *σ*(*s*) is also proportional to *s*). **C**. Mean response proportional to [log(*s*) + *c*]^2^ with Poisson (integer) response distribution, for which 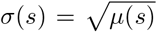. The panel on the right indicates the perceptual discrimination threshold (top) and the sensitivity (bottom) that arise from the calculation of Fisher Information, which are identical for all three representations.

First, Weber’s law can arise from a non-linear internal representational mean *µ*(*s*) (often referred to as a “transducer function”). If we assume that *µ*(*s*) is contaminated by additive Gaussian noise with variance *σ*^2^ [3, 33, 34]: *p*(*r*|*s*) ∼ 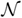[*µ*(*s*), *σ*], then 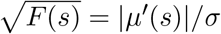 (see Methods). Thus, sensitivity to small stimulus perturbations is proportional to the derivative of the representational mean. Notice that this is a differential version of the standard measure of ‘d-prime’ in Signal Detection theory, which is used to quantify discriminability of two discrete stimuli (see Supplement). Under these conditions, sensitivity follows Weber’s law if the transducer is *µ*(*s*) ∝ log(*s*) + *c*, with *c* an arbitrary constant (Fig. 1A illustrates a case when *c* = 0, also see Methods). This logarithmic model of internal representation, due to Fechner [1, 5], is the most well-known explanation of Weber’s law.

Alternatively, a number of authors proposed that Weber’s law arises from representations in which noise amplitude grows in proportion to stimulus strength (sometimes called “multiplicative noise”) [3, 35–40]. Suppose representational mean *µ*(*s*) is proportional to stimulus strength (*s*), and is contaminated by Gaussian noise with standard deviation also proportional to *s*: *p*(*r*|*s*) ∼ 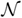 [*s, s*^2^]. The square root of FI for this representation again yields 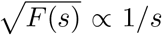, consistent with Weber’s law (see Methods). Note that unlike the previous case (in which Weber’s law arose from the nonlinear transducer), sensitivity in this case arises entirely from the stimulus-dependence of the noise variance (Fig. 1B).

Now consider a third case, inspired by neurobiology. Assume the stimulus is internally represented through neural spike counts that are Poisson-distributed with rate *µ*(*s*) (e.g. [41–43]). Despite the discrete nature of the spike count responses, FI may still be computed, and provides a bound on sensitivity. In this case, noise variance is equal to the mean response, and sensitivity is 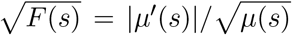, which gives rise to Weber’s law for a transducer function *µ*(*s*) ∝ [log(*s*) + *c*]^2^, where *c* is an integration constant (Fig. 1C, see Methods). Here, sensitivity reflects the combined signal-dependence of transducer and noise.

These three different examples demonstrate that an observation of Weber’s law sensitivity does not uniquely constrain an internal representation (see also [44–47]). In fact, these are three members of an infinite family of representations *p*(*r*|*s*) whose Fisher Information is consistent with Weber’s law. To make this non-identifiability problem more explicit, we introduce a simpler quantity which we dub *Fisher Sensitivity*, defined as:

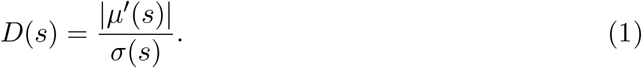

In general, Fisher Sensitivity provides a lower bound on the square root of FI [48] (see Methods), and is easier to compute, since it relies only on the first two moments of the response distribution. Its expression as a ratio of the change in response mean to standard deviation also provides an explicit connection to the “d-prime” measure used to quantify discriminability in Signal Detection theory (see Methods). For all three of the examples in the preceding paragraphs, this lower bound is exact (i.e., Fisher Sensitivity is identical to the square root of FI). But Fisher Sensitivity offers a direct and intuitive extension of the non-identifiability problem beyond these examples: To explain any measured pattern of sensitivity *D*(*s*), one can choose an arbitrary mean internal response *µ*(*s*) that increases monotonically and continuously, and pair it with an internal noise with variability *σ*(*s*) = |*µ*′(*s*)|*/D*(*s*). How can we resolve this ambiguity?

### Unified interpretation of power-law intensity percepts and Weber’s law sensitivity

The ambiguity described in the previous section can be resolved through additional measurements (or assumptions) of the mean or variance of internal representations, or the relationship between the two. In this section, we interpret perceptual magnitude ratings as a direct measurement of the representational mean, *µ*(*s*) [44, 49]. In a rating experiment, observers are asked to report perceived stimulus intensities by selecting a number from a rating scale (e.g. [7, 16, 17]). Suppose that these ratings reflect the observers’ internal response *r* (up to an arbitrary scale factor that depends on the numerical scale), and that averaging over many trials of *r* (drawn from *p*(*r*|*s*)) provides an estimate of the mean response, *µ*(*s*).

Using magnitude ratings, Stevens and others (e.g. [16, 50, 51]) showed that perceived intensity of many stimulus attributes can be well-approximated by a power law, *µ*(*s*) ∝ *s*^*α*^. The exponent *α* was found to vary widely across stimulus attributes ranging from strongly compressive (e.g., *α* = 0.33 for brightness of a small visual target) to strongly expansive (e.g., *α* = 3.5 for electric shock to fingertips). For stimulus attributes obeying Weber’s law, Stevens’ power law observations were interpreted as direct evidence against Fechner’s hypothesis of logarithmic transducers [11]. But the relationship of power law ratings to Weber’s law sensitivity was left unresolved. Over the intervening decades, magnitude rating measurements have generally been interpreted as arising from aspects of internal representation that are different from those underlying sensitivity (e.g. [12, 18, 21, 52]), or sometimes, measurements of magnitude ratings were dismissed altogether [9, 13].

Fisher Sensitivity offers a potential unification of power-law intensity percepts and Weber’s law sensitivity. First, we assume the observer whose discrimination behavior matches Weber’s law does so by optimally decoding an internal representation, achieving the Fisher Sensitivity: 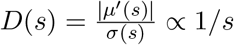. Substituting a power function, *µ*(*s*) = *s*^*α*^, and solving for *σ*(*s*) yields (Fig. 2A, see Methods):

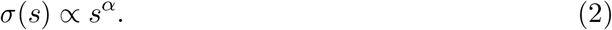

**Figure 2.**
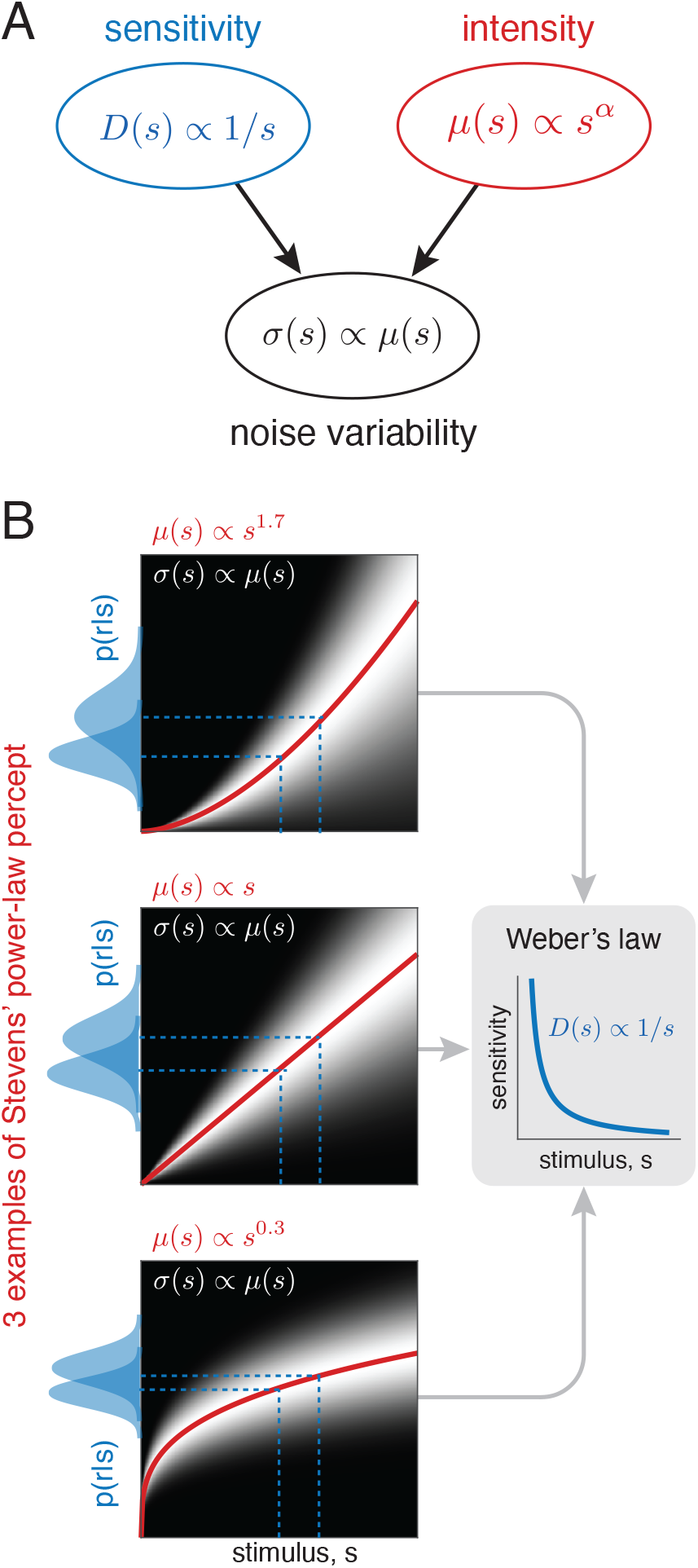
Unification of power-law intensity and Weber’s law sensitivity measurements. **A**. Using Fisher Sensitivity, perceptual sensitivity and intensity measurements can be combined to constrain the noise properties of an internal representation. In the particular case of Weber’s law, and power-law intensity ratings, this yields an internal representation with noise standard deviation proportional to mean response. **B**. This pattern of proportional internal noise serves to unify Weber’s law and power-law magnitudes for any exponent *α*, allowing for transducer functions that are expansive (*α >* 1, upper panel), linear (*α* = 1, middle), or compressive (*α <* 1, lower). Blue dashed lines indicate an example pair of stimuli that are equally discriminable in all three cases, as can be deduced qualitatively from the overlap of their corresponding response distributions (shown along left vertical edge of each plot, in shaded blue).

Thus, the standard deviation of the internal representation is proportional to its mean. This result holds for all values of *α*, and does not assume Gaussian internal noise, thus providing a generalization of the multiplicative noise example from the previous section (Fig. 1). Under these conditions, Weber’s law sensitivity can co-exist with a power-law intensity percept for *any* exponent (Fig. 2B).

### Connecting perceived intensity and discrimination of generalized intensity variables

The previous section provided a unification of three idealized relationships: Weber’s law for sensitivity, a power-law behavior for intensity ratings, and proportionality of mean and standard deviation of the internal representation. In this section, we consider generalizations beyond these relationships, and show that these can remain consistent under our framework.

Consider first the internal noise. Poisson neural noise implies a variance proportional to the mean spike count, a relationship that holds empirically for relatively low response levels [53]. At modest to high firing rates, spike count variance in individual neurons is generally super-Poisson, growing approximately as the square of mean response [53–55], consistent with the proportional noise assumption of the previous section. A modulated Poisson model has variance with both linear and quadratic terms, and can capture the relationship of spike count variance to mean response over the extended range [53, 54]:

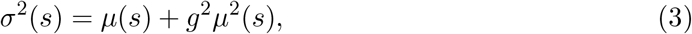

where the constant *g* represents the standard deviation of the modulation, and governs the transition from the Poisson range (smaller *µ*(*s*)) to the super-Poisson range (larger *µ*(*s*)) (Fig. 3A).

**Figure 3.**
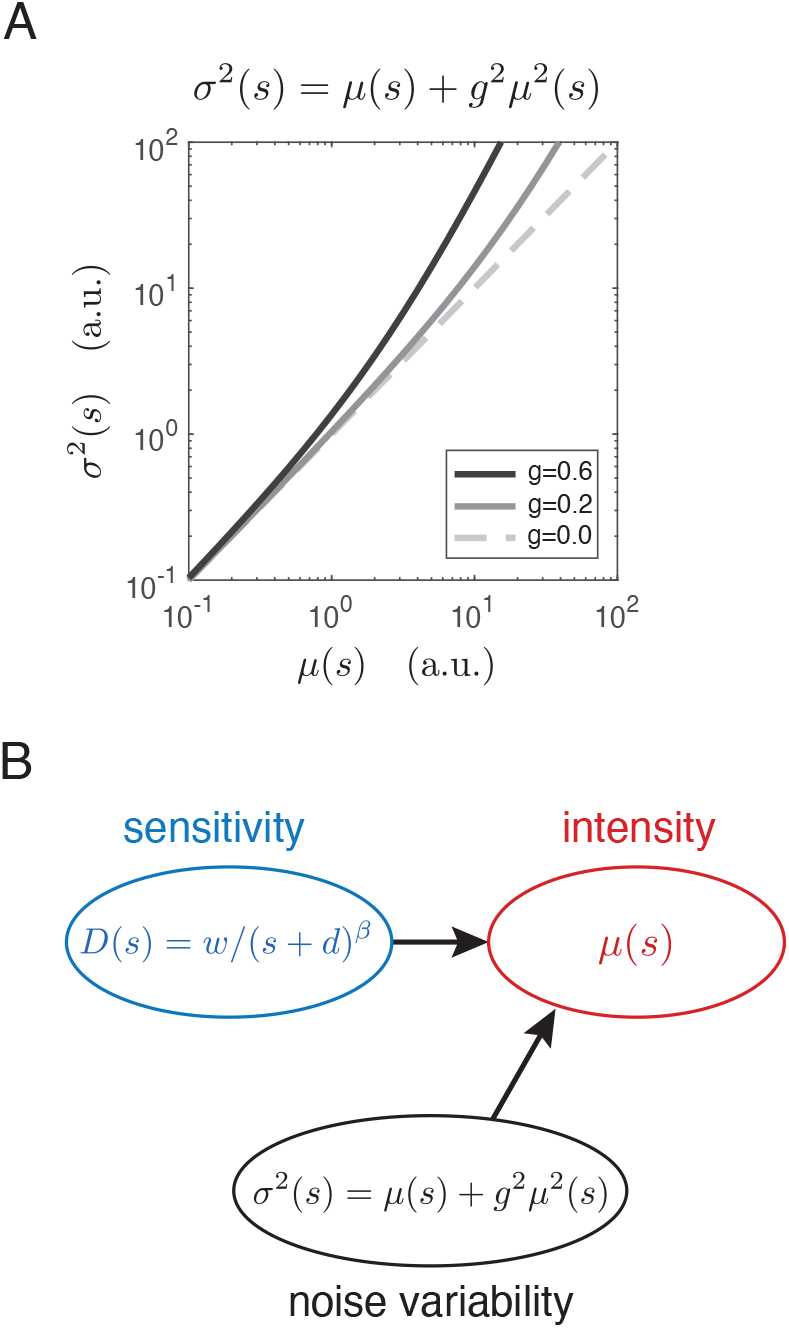
Generalization beyond the Weber range. **A.** Quadratic mean-variance relationship for a modulated Poisson model of sensory neurons [53, 55]. Behavior is Poisson-like at low intensities (i.e., when *µ*(*s*) is much less than 1*/g*^2^, then *σ*^2^(*s*) ∼ *µ*(*s*)), and super-Poisson at higher intensities (when *µ*(*s*) is much greater than 1*/g*^2^, then *σ*^2^(*s*) ∼ *µ*^2^(*s*)). **B**. Using Fisher Sensitivity, a generalized form of Weber sensitivity can be combined with the mean-variance relationship in panel A to generate numerical predictions of perceived stimulus intensity *µ*(*s*) (see examples in Fig. 4).

Perceptually, both Weber’s law for sensitivity and the power-law for perceptual magnitudes are known to fail, especially at low intensities (e.g. [7, 56]). A generalized form of Weber’s law (e.g. [57]) has been proposed to capture sensitivity data over broader range of intensity:

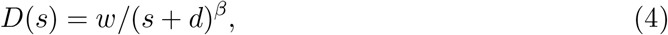

where *d* is a constant that governs sensitivity at low intensities, the exponent *β* determines deviation from Weber’s law at high intensities, and *w* is a non-negative scaling factor. Weber’s law corresponds to the special case of *d* = 0 and *β* = 1.

To test the generalization of our unified framework, we used Fisher Sensitivity to combine the modulated Poisson noise model (Eq. (3)) with fitted versions of this generalized form of Weber’s law (Eq. (4)), and to generate predictions of *µ*(*s*) (illustrated in Fig. 3B). We then compared these predictions to averaged perceptual intensity ratings. The predictions rely on the choice of three parameters: *g* that determines the transition from Poisson to super-Poisson noise, an integration constant *c*, and a scale factor that accounts for the range of the rating scale used in the experiment (see Methods). We examined predictions for five different stimulus attributes, for which both sensitivity and rating scale data (averaged across trials) are available over a large range of stimulus intensities. Fig. 4 shows results for: 1) concentration of sucrose [“sweetness”, [58, 59]]; 2) concentration of sodium chloride [“saltiness”, [58, 59]]; 3) amplitude of white noise [auditory loudness, [60, 61]]; 4) amplitude of 1000 Hz pure tone [auditory loudness, [61, 62]]; and 5) amplitude of a sinusoidal grating [visual contrast, [51, 57]].

**Figure 4.**
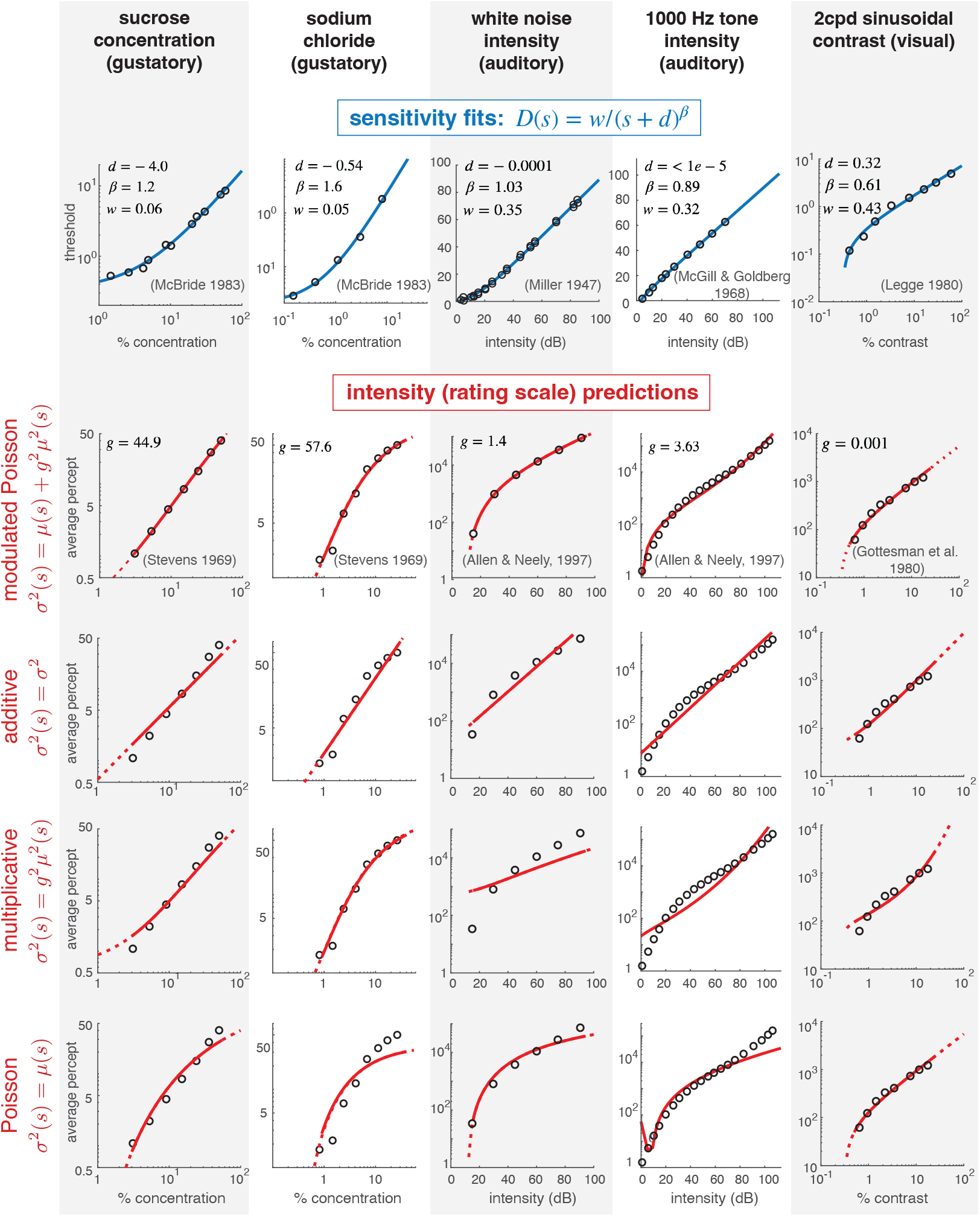
Predictions of perceived intensity from sensitivity, for five different sensory attributes. Top row: For each attribute, we fit a three-parameter generalized form of Weber’s law (Eq. (4), blue curves) to measured discrimination thresholds (hollow points). Optimal parameter values for each attribute are indicated. Bottom four rows: Fitted sensitivity functions are equated to the Fisher Sensitivity relationship (Eq. (1)), assuming one of four different mean-variance relationships (equations, left side), to generate predictions of perceived intensity *µ*(*s*) (red curves). The modulated Poisson and multiplicative noise predictions depend on parameter *g*, and the additive noise prediction depends on parameter *σ*^2^. In addition, all predictions depend on an integration constant *c* and an overall multiplicative scale factor (see Methods). All parameters are adjusted to best fit average perceptual rating scale measurements (hollow points).

The sensitivity curves vary substantially across these stimulus attributes, but all are well-fit by the generalized Weber functional form (blue curves, first row of Fig. 4). In all cases, the rating scale data are well-predicted by combining the sensitivity fit with the modulated Poisson noise model of Eq. (3) (red curves, second row, Fig. 4). Moreover, we find that reduction to simpler noise models (multiplicative, or Poisson) that are special cases of the full model provide worse predictions for many cases (rows 4 and 5, Fig. 4). Specifically, when *g* is small (as in the case of visual contrast), the modulated Poisson model behaves similarly to a standard Poisson model, but the multiplicative model fit is poor. When *g* is large (as in the case of tasting sodium chloride), the noise model behaves similarly to the multiplicative noise model, but the Poisson model fit is poor. Note that the standard Poisson model has one less parameter than the other models.

The additive noise model is also worse than the modulated Poisson model, but generally outperforms the other two (Fig.4, row 3). In the five stimulus domains examined, we did not observe any systematic pattern of model parameters across stimulus categories (for either the sensitivity fit or the rating scale predictions). But examination of additional stimulus domains using this type of concurrent measurement may reveal such patterns.

## Discussion

Stimulus magnitude and sensitivity are amongst the most widely assessed perceptual characteristics [63, 64], but the relationship between the two has proven elusive. In this article, we’ve proposed a framework that relates these characteristics to two fundamental properties of internal representation – a nonlinear “transducer” that expresses the mapping of stimulus magnitude to the mean internal representation, and the stimulus-dependent amplitude of internal noise. Our proposal relies on two assumptions that link perceptual measurements to these properties: (1) sensitivity (the inverse of the discrimination threshold) reflects a combination of the transducer and the noise amplitude, as expressed by Fisher Sensitivity; and (2) absolute judgements (specifically, those obtained through average ratings of stimulus intensity) reflect the value of the transducer. This combination allows a unified interpretation in which intensity and sensitivity reflect a single underlying representation, providing a potential link to physiology.

Our framework relies on several assumptions. First, we restrict ourselves to continous scalar stimulus domain, and an internal representation that is differentiable with respect to the stimulus (so that Fisher Information is well-defined). Throughout, we rely on Fisher Sensitivity, an intuitive and tractable lower bound on the square root of Fisher Information. The two are equivalent for the Weber’s law examples shown in Figs. 1 and 2, but not for the data fitting examples of Fig. 4 (in the Supplement, we provide an additional example in which the two quantities differ). We assume human perceptual sensitivity achieves (or is at least proportional to) the Fisher Sensitivity bound. More specifically, we assume that human responses in a perceptual discrimination task reflect optimal extraction of information from a noisy internal representation, as suggested by a number of studies linking physiology to perception (e.g., [65–69]). Finally, we assume that absolute intensity judgements reflect a transducer function that corresponds to the mean of the internal representation.

To develop and test this framework, we have focused on attributes that obey Weber’s law, and its modest generalizations. Despite its ubiquity, the relationship between Weber’s law and the underlying representation has been contentious. In the late 19th century, Fechner proposed that perceptual intensities correspond to integrated sensitivity [1], and in particular predicted that Weber’s law sensitivity implied a logarithmic internal representation. Using rating scales as a form of measurement, Stevens instead reported that many sensory variables appeared to obey a power law, with exponents differing substantially for different attributes [11]. Stevens interpreted this as a refutation of Fechner’s logarithmic transducer. In order to explain the discrepancy between Fechner and Stevens’ proposals, a number of authors suggest that perceptual intensities and sensitivity reflect different stages of processing, bridged by an additional nonlinear transform. Specifically, [18] proposes a type of sensory adaptation, [19] reflects additional sensory processing, and [22] incorporates an additional cognitive process. Our framework offers a parsimonious resolution of these discrepancies, by postulating that perceptual intensity and sensitivity arise from different combinations of the mean and variance of a common internal representation.

It is worth noting that while Fechner’s integration hypothesis is inconsistent with Stevens’ power law measurements, it appears to be consistent with many supra-threshold intensity measurements. Specifically, experimental procedures involving supra-threshold comparative judgements (e.g. maximum likelihood difference scaling methods, categorical scales and bisection procedures [17, 40, 58, 70]) seem to reflect integration of sensitivity, whereas experimental procedures that require absolute judgements (e.g. rating scales [17, 51, 71]) yield different functions that we’ve interpreted as reflecting the mean of internal representation. In the case of Weber’s law, the integrated sensitivity is logarithmic, consistent with Fechner’s interpretations, *regardless* of the underlying transducer-noise combination (e.g., Fig. 1)! Under this interpretation, our framework can provide a natural unification of Stevens’ power law magnitude ratings, Weber’s law sensitivity, and Fechner’s logarithmic supra-threshold distances (Fig. 5). Further empirical studies will be needed to verify or refute these relationships.

**Figure 5.**
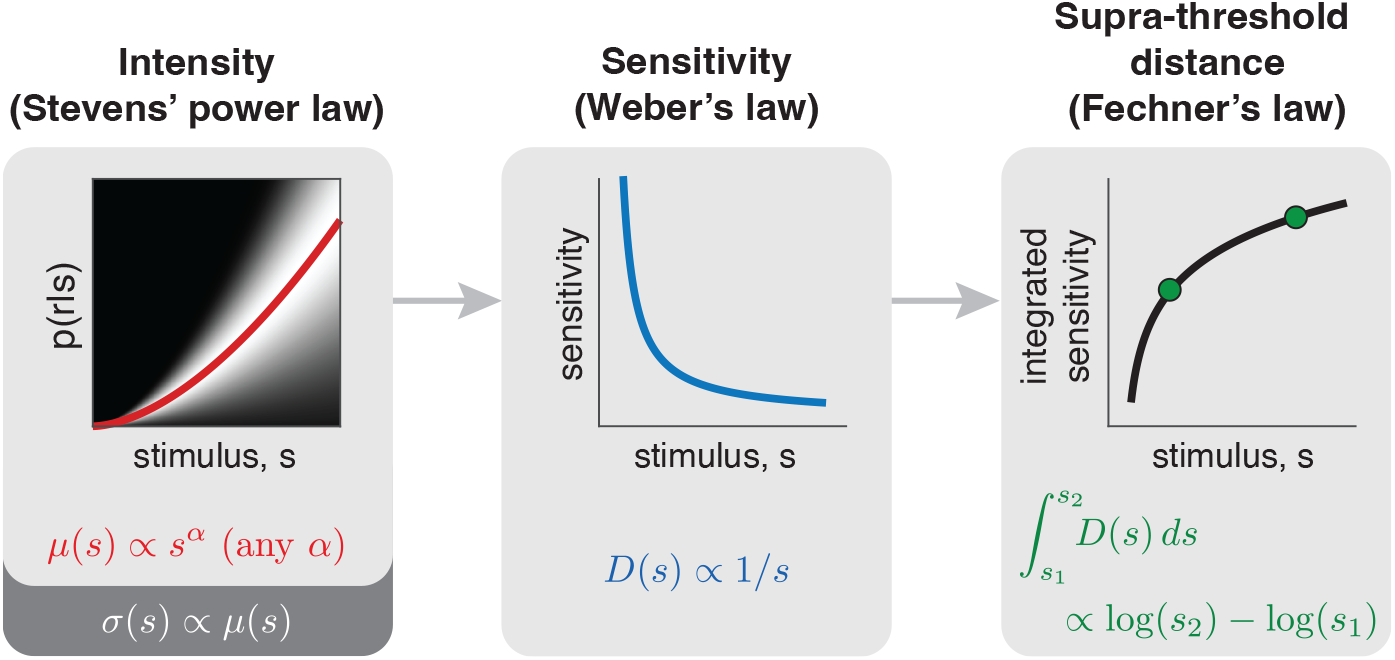
Extension of the Fisher Sensitivity framework to supra-threshold perceptual distances. Weber’s law is consistent with Stevens’ power law (for any exponent, *α*) as long as the standard deviation of the noise scales with the same exponent (left and middle panels; see also Fig. 2B). In addition, under the assumption that perceived supra-threshold distances correspond to *integrated* sensitivity, these will correspond to differences in logarithmically mapped stimuli, providing a modified interpretation of Fechner’s law. Under these conditions, all three “laws” co-exist in a consistent framework, each describing measurements that access different aspects of a common underlying representation.

This subtle distinction between comparative and absolute judgement is at the heart of multiple debates in perceptual literature. For example, it arises in discussions of whether perceptual noise is additive or multiplicative in visual contrast (e.g. [40, 44, 72]). We have proposed that mean and variance of internal representations can be identified through the combination of absolute and discriminative judgements, because the two measurements reflect different aspects of the representation. On the other hand, if supra-threshold comparative judgements reflect *integrated* local sensitivity, they will not provide additional constraints on internal representation beyond threshold sensitivity measurements, and combining these two measurements cannot resolve the identifiability issue. This provides, for example, a consistent interpretation of the analysis in [40], which shares the logic of our approach in seeking an additional measurement to resolve non-identifiability of sensitivity measurements, but reaches a different conclusion regarding consistency of additive noise.

Several other theoretical or experimental constraints have been proposed to resolve the identifiability issue, including imposing a common criterion between two discrimination tasks [72], connecting the response accuracy for the first and the second response in a four-alternative choice [31], and connecting discrimination to an identification task [47]. An open question is whether our framework can be extended to account for these more diverse perceptual scenarios.

Our examination of the particular combination of Weber’s law sensitivity with power-law intensity percepts led to the conclusion that the standard deviation of internal noise in these cases should vary in proportion to the mean response. While such “multiplicative noise” has been previously proposed as an explanation for Weber’s law [3, 35–37], it has generally been described in the context of a linear transducer (as in Fig. 1). In our framework, we find that this form of noise (standard deviation proportional to the mean) is sufficient to unify Weber’s and Stevens’ observations for the complete family of power-law transducers, regardless of exponent. An additional prediction of this model is that the standard deviation of perceptual magnitude ratings should grow proportionally to the mean rating (consistent with Fig. 2B). This is consistent with findings of a number of previous studies (e.g. [10, 73, 74]). For example, Green and Luce showed that when observers were asked to rate 1000 Hz tone loudness, their coefficient of variations (standard deviation divided by the mean) in the ratings are near-constant for a wide range of intensities [73].

The proportionality of the mean and standard deviation of a stimulus representation offers a potential interpretation in terms of underlying physiology of neural responses. We con-sidered, in particular, recently proposed “modulated Poisson” models for neural response which yields noise whose variance grows as a second-order polynomial of the mean response [53, 54, 75]. The noise of the summed response over a population of such neurons would have the same structure (see Supplement). At high levels of response, this allows a unification of Weber’s law and Stevens’ power law. At lower levels, it produces systematic deviations that lead to consistent predictions of ratings for a number of examples (Fig. 4). Recent generalizations of the modulated Poisson model may allow further refinement of the perceptual predictions [76]. For example, at very low levels of response, sensory neurons exhibit spontaneous levels of activity that are independent of stimulus drive [34], suggesting that inclusion of an additive constant in Eq. (3) could improve predictions of perceptual detection thresholds [77].

We’ve restricted our examples to perceptual intensity attributes that obey Weber’s law, but the proposed framework is more general. In particular, the Fisher Information bound holds for any noisy representation, and has, for example, been applied to the representation of sensory variables in neural population responses [25, 28, 29]. In some cases, these attributes exhibit Weber’s law behavior, which may be ascribed to the combination of heterogeneous arrangements of neural tuning curves along with noise properties of individual neurons [78– 80]. For example, neurons in visual area MT selective for different speeds have tuning curves that are (approximately) shifted on a logarithmic speed axis [81]. Under these conditions, an independent response noise model yields Fisher Information consistent with Weber’s law [82, 83]. More generally, changes in a stimulus attribute may cause changes in both the amplitude and the pattern of neuronal responses, which, when coupled with properties of internal noise, yield predictions of sensitivity through Fisher Information. Specifically, the abstract internal representation that we have assumed for each perceptual attribute corresponds to the projection of high-dimensional noisy neuronal responses onto a decision axis for perceptual judgements (e.g. [54, 84, 85]). Although discrimination judgements for a stimulus attribute are generally insufficient to uniquely constrain underlying high-dimensional neuronal responses, the one-dimensional projection of these responses provides an abstract but useful form for unifying the perceptual measurements.

Our framework enables the unification of two fundamental forms of perceptual measurement – magnitude judgement and sensitivity – with respect to a common internal representation. However, the study of perception is diverse and mature, with numerous additional perceptual measurements [86] whose connection to this framework could be explored. The descriptive framework outlined here also raises fundamental questions about the relationship between internal representation mean and noise. The forms of both noise and transducer may well be constrained by their construction from biological elements, but may also be co-adapted to satisfy normative goals of efficient transmission of environmental information under constraints of finite coding resources [87–89]. Exploration of these relationships provides an enticing direction for future investigation.

## Methods

### Fisher Information

For a stimulus attribute *s*, the Fisher Information (FI) is derived from the conditional distribution of responses given the stimulus, *p*(*r*|*s*), and expresses the relative change in response distribution when the stimulus *s* is perturbed:

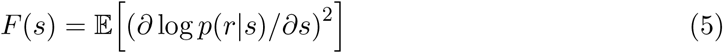

where the expectation is taken over the distribution *p*(*r*|*s*)[90]. Intuitively, Fisher information converts a description of the internal noisy representation, *p*(*r*|*s*), into a measure of the precision (inverse variance) with which the stimulus is represented [91]. The definition relies only on the differentiability of the response distribution with respect to *s* and some modest regularity conditions [91], but does not make assumptions regarding the form of the noisy response distribution. Either *s* or *r* can be vector-valued, but for our purposes in this article, we assume a one-dimensional stimulus attribute, and thus the internal representation *r* that is relevant to the discrimination experiment is also effectively one-dimensional.

In statistics and engineering communities, FI is often used in the context of the “Cramér-Rao bound”, an upper bound on the precision (inverse variance) attainable by an unbiased estimator [91]. It was first proposed as a means of quantifying perceptual discrimination by Paradiso [29], and further elaborated for neural populations by Seung and Sompolinsky [25]. In this context, the square root of Fisher Information provides a bound on perceptual precision (sensitivity) [30], and may be viewed as a generalization of “d-prime” [27], the traditional metric of signal detection used in psychophysical studies [3] (see Supplement).

### Three example representations yielding Weber’s law sensitivity

The three example representations shown in Fig. 1 are each consistent with Weber’s Law, but differ markedly in their response distributions. Below, we derive each of these.

### Additive Gaussian noise

Assume the internal representation has mean response *µ*(*s*), and is contaminated with additive Gaussian noise with standard deviation *σ*:

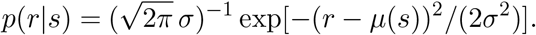

Substituting into Eq. (5) and simplifying yields 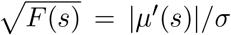. Weber’s Law corresponds to sensitivity proportional to 1*/s*, and thus we require a transducer such that |*µ*′(*s*)| ∝ 1*/s*. If we assume monotonicity, the transducer is uniquely determined (up to an integration constant and a proportionality factor) via integration: *µ*(*s*) ∝ log(*s*) + *c*.

### “Multiplicative” Gaussian noise

Assume a representation with identity transducer *µ*(*s*) = *s* and Gaussian noise such that the amplitude scales with the mean 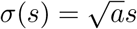:

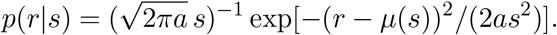

Substituting into Eq. (5) and simplifying again yields Weber’s Law: 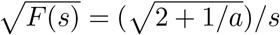.

### Poisson noise

Assume the internal response *r* is an (integer) spike count, drawn from an inhomogeneous Poisson process with rate *µ*(*s*), a widely-used statistical description of neuronal spiking variability. Then

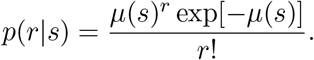

In this case, 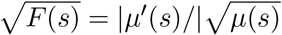. Assuming Weber’s law, we can again derive the form of the transducer: *µ*(*s*) ∝ [log(*s*) + *c*]^2^ for some constant *c*.

### Fisher Sensitivity

In general, Fisher Information can be difficult to compute and often cannot be expressed in closed form. A lower bound for the square-root of Fisher Information, which we term *Fisher Sensitivity*, is more easily computed and interpreted, because it depends only on the mean and variance of the distribution. Specifically, we define Fisher Sensitivity as:

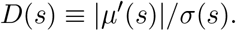

Its role as a lower bound can be derived using the Cauchy-Schwartz inequality for continuous density *p*(*x*)*dx*:

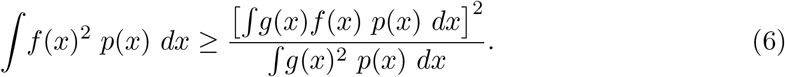

Making the following substitutions:

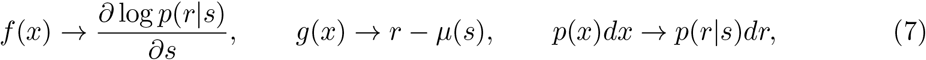

the left side of Eq. (6) is equal to the Fisher Information (defined in Eq. (5)), and the right side is equal to the squared Fisher Sensitivity:

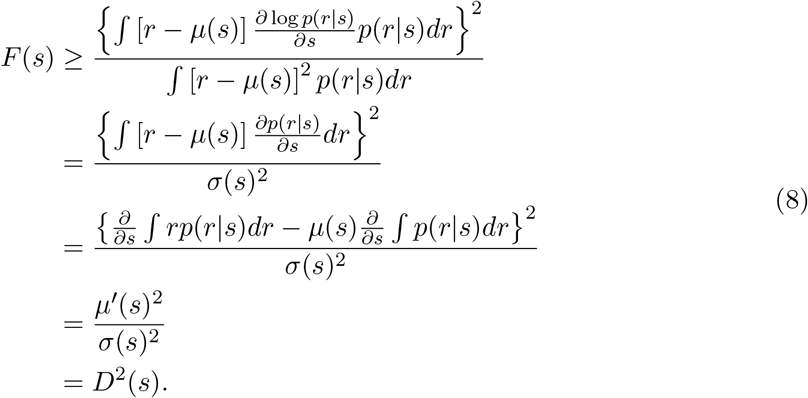

Fisher Sensitivity generalizes to multi-dimensional response vectors (e.g., a neural population), by replacing the inverse variance with the Fisher Information matrix, and projecting this onto the gradient of the mean response [92, 93]. The derivation of the full bound for the multi-dimensional case (both stimuli and responses) may be found in [48].

In the examples of Fig. 1 and Fig. 2, the lower bound is exact: Fisher Sensitivity is equal to the square-root of Fisher Information. An equivalent expression for Fisher Sensitivity has also been derived by assuming a minimal-variance unbiased linear decoder [94]. Compared to our interpretation as a lower bound, this interpretation has the advantage of being an exact expression of Fisher Information, but the disadvantage of relying on restrictive decoding assumptions.

### Relationship of Fisher Sensitivity to Signal Detection theory

In Signal Detection theory, discriminability between two stimulus levels *s*_1_ and *s*_2_ is typically summarized using the measure known as “d-prime”. To relate this to Fisher Sensitivity, we assume a simple form sometimes used in the perception literature:

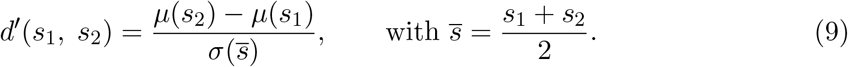

Assuming *s*_1_ and *s*_2_ are two values on a continuum, and that *µ*(*s*) is differentiable, we can express the two internal responses using a first-order (linear) Taylor approximation:

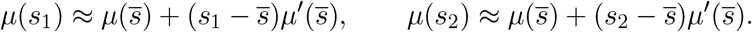

Substituting these into Eq. (9) gives:

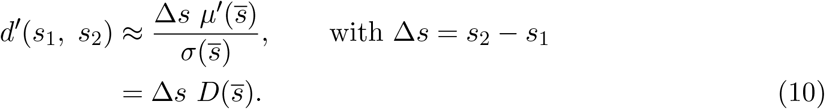

That is, Fisher Sensitivity expresses the slope at which d-prime increases with stimulus separation. Setting d-prime equal to a criterion level *d*^*^ and solving for the stimulus discrimination threshold gives:

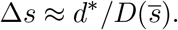

That is, discrimination thresholds are inversely proportional to Fisher Sensitivity. This relationship was used to fit the data for Fig. 4.

### Internal representations consistent with Weber’s law and Stevens’ Power law

Using Fisher Sensitivity and assuming monotonicity of *µ*(*s*), Weber’s law can be expressed as: 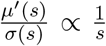. To identify *µ*(*s*) and *σ*(*s*), we combine this with magnitude ratings, which we assume provide a direct measurement of *µ*(*s*). Assume the magnitude ratings follow a power law [16]. Then *µ*(*s*) ∝ *s*^*α*^, with derivative *µ*′(*s*) = *αs*^*α*− 1^. Substituting into the equation for Weber’s law and solving gives *σ*(*s*) ∝ *s*^*α*^. That is, Weber’s law can arise when both *µ*(*s*) and *σ*(*s*) follow a power law with the same exponent, *α*. Note that this result holds for all exponents.

### Data Fitting

To examine the validity of our framework beyond Weber’s range, we analyzed five different sensory attributes (Fig. 4). For each, we first fit a generalized form of Weber’s Law [20] to perceptual sensitivity data:

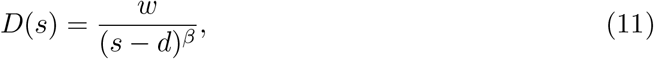

in which *d* is an unrestricted additive constant, *β* is a non-negative exponent, and *w* is a non-negative scaling factor. These three parameters were optimized to minimize squared error of the measured thresholds (inverse sensitivity).

Next, we combined the fitted sensitivity model with a model of internal noise to generate a prediction for the mean percept, *µ*(*s*), which was then compared with rating measurements. This was carried out for four different noise models: modulated Poisson, additive, multiplicative, and Poisson (corresponding to bottom four rows of Fig. 4, respectively). We derive the corresponding expressions for *µ*(*s*) below.

### Modulated Poisson noise

Our primary predictions assume a modulated Poisson noise model [53] with mean-variance relationship:

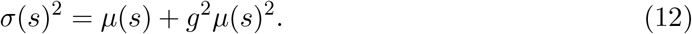

The transducer *µ*(*s*) is obtained by solving the differential equation that arises by substituting this variance expression into the Fisher Sensitivity of Eq. (1), and equating this with the generalized form of Weber’s law (Eq. (11)):

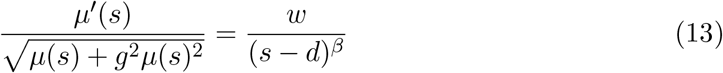

The solution may be expressed in closed form:

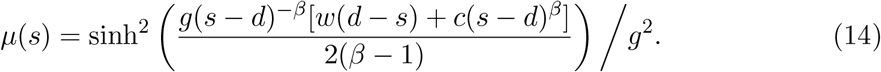

The parameters {*d, β, w*} are constrained to values obtained when fitting the sensitivity data, and three remaining parameters are adjusted to minimize squared error with the log-transformed rating data. The first is *g*, which governs the transition from Poisson to super-Poisson noise behavior (large *g* indicates an early transition). The second is *c*, an integration constant that arises from solving the differential equation for *µ*(*s*). The last parameter is an overall scale factor (not indicated), which rescales the predicted intensity values to the numerical range used in the associated rating experiment.

### Additive Gaussian noise

As for the full modulated Poisson model, we first fit the generalized Weber’s law to discrimination data, and locked the parameters {*d, β, w*}. Then we solve a differential equation arising from equating Fisher Sensitivity with the generalized Weber’s Law:

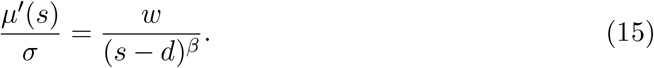

The solution for *µ*(*s*) in this case may also be expressed in closed form:

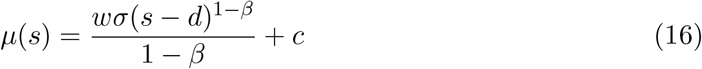

The integration constant *c*, the constant *σ*, and an overall scaling factor are adjusted to fit *µ*(*s*) to the rating data (minimizing the squared error between logarithmically transformed rating data and the function).

### Poisson noise

Following a similar procedure for the case of additive Gaussian noise, we find a closed-form solution for *µ*(*s*) using Poisson noise and Fisher Sensitivity:

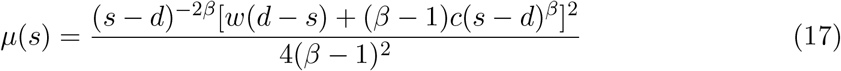

Again, the integration constant *c* and overall scaling factor are optimized to fit the rating data.

### Generalized multiplicative noise

Here, we assume a noise mean-variance relationship 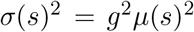, which is the choice that enables the co-existance of the classic form of Weber’s law and Stevens’ power law. As in previous cases, we substitute this into the expression for Fisher Sensitivity to obtain a prediction for *µ*(*s*):

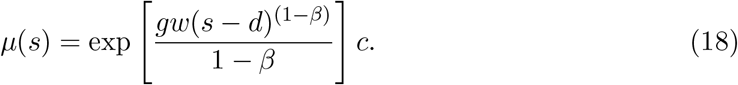

Note that, as for the full noise model of Eq. (12), comparison to the rating data involves estimation of three parameters: the noise parameter *g*, an integration constant *c*, and a scaling factor.

## Acknowledgments

We thank Nikhil Parthasarathy, Mike Landy, Tony Movshon, David Brainard, Robbe Goris, Bill Geisler, Larry Maloney, Jacob Cheeseman and members of the Center for Computational Neuroscience at the Flatiron Institute for helpful discussions and suggestions. In addition, we are grateful to the editor and three reviewers for their constructive questions and suggestions, which encouraged a variety of improvements in the manuscript.

## Appendices

### Fisher Sensitivity and Fisher Information for Gaussian responses

Consider the general case of an internal representation with Gaussian noise having stimulus-dependent mean and variance:

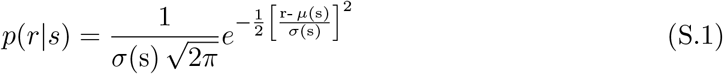

The Fisher Information of this representation can be computed as:

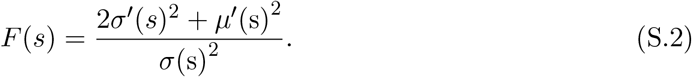

For some cases, this is equal to the squared Fisher Sensitivity. Specifically, for the constant-variance case (additive noise, top panel of Fig. 1), *σ*′(*s*) = 0, and *F* (*s*) = *µ*′(*s*)^2^*/σ*(*s*)^2^. Also, when *σ*(*s*) ∝ *µ*(*s*) (e.g., Fig. 2), then *F* (*s*) ∝ *µ*′(*s*)^2^*/σ*(*s*)^2^. But in general, these two quantities are different.

To examine how close Fisher Sensitivity is to the square-root of Fisher Information in the Gaussian case, we can write the Gaussian standard deviation *σ*(*s*) as a function of the mean: *σ*(*s*) = *h*[*µ*(*s*)]. Then Eq. (S.2) can be re-expressed as the following:

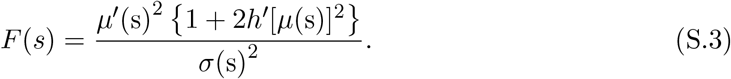

In general when *h*[·] is not a constant, if the standard deviation *σ*(*s*) = *h*[*µ*(*s*)] varies slowly as a function of the stimulus (or when *h*′[*µ*(*s*)] is small), the lower bound is relatively tight.

## Notes

### Competing Interest Statement

The authors have declared no competing interest.

### Summary of Updates

Copy editing for the final publication.

